# Holistic assessment of large old trees: a framework and its application for Romania

**DOI:** 10.1101/2020.03.24.004812

**Authors:** Viorel Arghius, Cristian Malos, Vlad Macicasan, Tibor Hartel

**Affiliations:** Faculty of Environmental Science and Engineering, Babes-Bolyai University, Cluj-Napoca, 30 Fântânele St., Romania; Hungarian Department of Biology and Ecology and Center of Systems Biology, Biodiversity and Bioresources, Babes-Bolyai University, Cluj-Napoca, Romania

**Keywords:** large old trees, natural values, cultural values, conservation regulations, Romania

## Abstract

Large old trees are keystone ecological structures and have exceptional sociocultural values. Still, holistic approaches to national assessments of large old trees are scarse in the scientific literature. Here we propose and apply a holistic framework to understand the distribution of large old trees, the formal regulations targeting the conservation of large old trees and the ways large old trees are present in the popular news in Romania. There were 4032 large old tree records in Romania most of the records being concentrated in the Central and North-Western part of Romania. The number of tree records decreases with the decreasing terrain accessibility. Almost 50% of the large old tree records are in areas not covered by nature conservation regulations and 2/3 of the terrestrial protected areas have no open access large old tree records, so far. We identified five formal regulations which could be relevant for large old tree conservation in Romania, however, only two of these explicitly targets large old trees. The lack of knowledge and interest, the lack of expertize, institutional capacity, vested interests (corruption) and inconsistencies within the regulations are the major barriers in the formal protection of large old trees. We also identified several opportunities for the local initiatives to protect large old trees, based on the current legislative frameworks. We identified 259 internet news targeting large old trees in Romania which reveals a wide range of values and concerns associated to large old trees at the level of the local communities. While discussing our results we highlight the benefits of a wider adoption of our approach for science, decision making and local initiatives to save large old trees.

## Introduction

A holistic approach which combines ecological, social and institutional dimensions has a great potential to leverage effective biological conservation strategies in human shaped landscapes (Abson et al., 2016; Hanspach et al., 2014). Focusing in ecosystem structures which acts as ecological keystones (i.e. have disproportionately large value for biodiversity and ecosystem function) and are flagships (i.e. are recognized as charismatic and important at the level of the human society) can help prioritizing nature conservation actions (Lindenmayer et al., 2015). Large old trees have several features which provides them exceptional ecological and sociocultural values including large cavities in the trunk, their hollowing and dead parts (Lindenmayer and Laurance, 2016) and their long age, which provides them cultural and historical legacy value for the local communities (Blicharska and Mikusiński, 2014; Lindenmayer, 2016). Large old trees are often present in human shaped environments and their national assessment provides opportunities to understand and share with the broad academic and policy community their status, national level regulations and the local initiatives which aims to save, popularize or protect these trees. Nevertheless large old trees often induces negative perceptions and attitudes in people and often have no direct monetary value (such as the timber trees), resulting in their removal despite the acknowledgement of their multiple socio-cultural values (Hartel et al., 2017; Lindenmayer, 2016; Orłowski and Nowak, 2007). Furthermore large old trees are often ignored (or are only indirectly addressed) by environmental and conservation policies at international and national levels (Lindenmayer et al., 2014). As a consequence their maintenance and protection largely relies on the ways how the existing nature conservation and environmental legislation is interpreted and the intent existing at the level of formal and informal institutions which are covering the tree.

Despite the lack of their formal recognition in the influential environmental and nature conservation strategies and regulations, several European countries started to inventory their large old trees and open databases exists which makes the informations about large old trees available to the broad public. Examples includes Sweden (Östberg et al., 2018; https://tradportalen.se/), Hungary (http://www.dendromania.hu/index.php), UK (http://www.ancienttreeforum.co.uk/), Romania (https://arboriremarcabili.ro/en/about-project/) or the international platform ‘Monumental Trees’ (https://www.monumentaltrees.com/en/records/nor/). Furthermore recently there is an increase in the academic papers presenting the regional or national status of large old trees, such as Cyprus (Anestiadou et al., 2017), Czech Republic (Uradníček et al., 2017), Italy – (Asciuto et al., 2015; Marco and Alessandro, 2008), Poland (Orłowski and Nowak, 2007; Tomas and Dorota, 2009) and Turkey (Polat, 2017). Large old trees are ideal targets for citizen science type of data collection in some countries. For example in the United Kingdom and Sweden tens of thousands of large old tree are recorded with the participation of a large number of volunteers (Butler, 2014, Östberg et al., 2018; Trädportalen – https://tradportalen.se/).

While valuable informations about the extent and environmental determinants of the large old tree are increasingly available recently (see above), holistic approaches which integrates the national level informations on large old trees, formal regulations and local initiatives to protect these trees are still virtually lacking. The overarching goal of this study is to provide comprehensive synthesis of the known large old trees of Romania, present the regulatory context within which large old trees are or can be protected and provide a glimpse in the local initiatives for protecting large old trees, based on the assessment of the electronic mass media (internet news). Knowing the current knowledge on large old trees helps prioritizing future research activities. Understanding the national level regulatory context for large old tree conservation can help in identifying the challenges for improving the regulations but also on identifying the opportunities to protect large old trees within the existing legislation context. Nevertheless, understanding the ways how the large old trees are present in the electronic news can help identifying the societal values, concerns and initiatives related to and targeting these trees. All three dimensions together can largely improve the knowledge on large old trees as well as their conservation opportunities at local, regional and national scales.

The first inventory of large old trees as natural monuments in Romania was carried out in 1925, the inventory targeting especially the trees with a great historical and cultural value (Mohan et al., 1993a). Since then, references on large old trees have been made in various publications (e.g. Pop and Sălăgeanu, 1965, Giurescu, 1975, Mohan et al., 1993, Bolea et.al., 2013a, Popa, 2016, Nădi□an and Bârda, 2016, Bolea, 2017, Moga et al., 2016, Hartel et al., 2018, Vasile et al., 2019). This is the first holistic systematic knowledge review on the large old trees from Romania. Specifically this study have the following objectives (O):

*(O1)* To present the current knowledge on the distribution of large old trees in Romania.
*(O2)* To overview the formal policies with relevance for the protection of large old trees in Romania.
*(O3)* To understand the socio-cultural values of large old trees through internet news content analysis.

In the following we define large old trees and then present a framework we used to collect the relevant informations in order to achieve our objectives. This will be followed by methodology description and the presentation of results. While discussing our results, we provide future directions for large old trees conservation in Romania with the following dimensions: research, civil society (citizen science, local level large old tree conservation) and policy.

### Defining large old trees

A tree can be defined as ‘large old’ based on several contextual criteria such as the ecosystem type and region where the tree developed, the biological characteristics of the tree species (e.g. age) and subjective judgements of the authors (Lindenmayer and Laurance, 2016). The trunk circumference (Circumference at Breast Height, CBH), or diameter (DBH) of the mature trees at *cca* 130 cm (sometimes 150 cm) height is often used as subjective indicator for considering a tree ‘large old’ (Lindenmayer and Laurance, 2016; Úradníček et.al., 2017; Hartel et al., 2018). While it is recognized that large trunk circumference may not always indicate old age it is assumed that within the same environmental contexts and tree species, trees with ticker trunk are older than the smaller trees. Besides the remarkably thick trunk, large old trees often have aged bark which is sometimes detaching, prominent hollows, dried parts and are rich in biodiversity (Lindenmayer and Laurance, 2016). Table 1 presents a taxa specific lower threshold size of trunk circumferences for considering a mature tree to be ‘large old’ for the existing Romanian tree records. The threshold values were established both by considering the available guidelines (e.g. Practical guidance published by the Woodland Trust, http://www.woodlandtrust.org.uk) as well as our collective understanding of large old trees in Romania, based on intense tree inventory and old tree documentations in the past years (see e.g. Hartel et al., 2013; Hartel et al., 2018; Remarkable Trees of Romania, 2020, www.arboriremarcabili.ro).

**Table 1.**
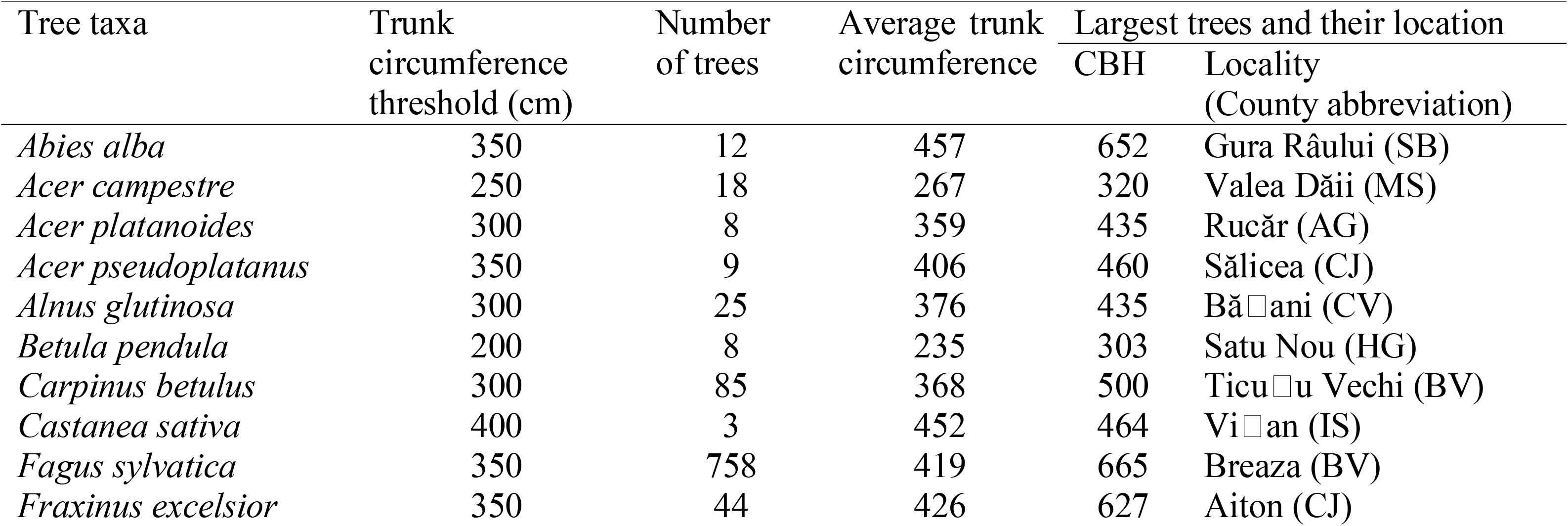
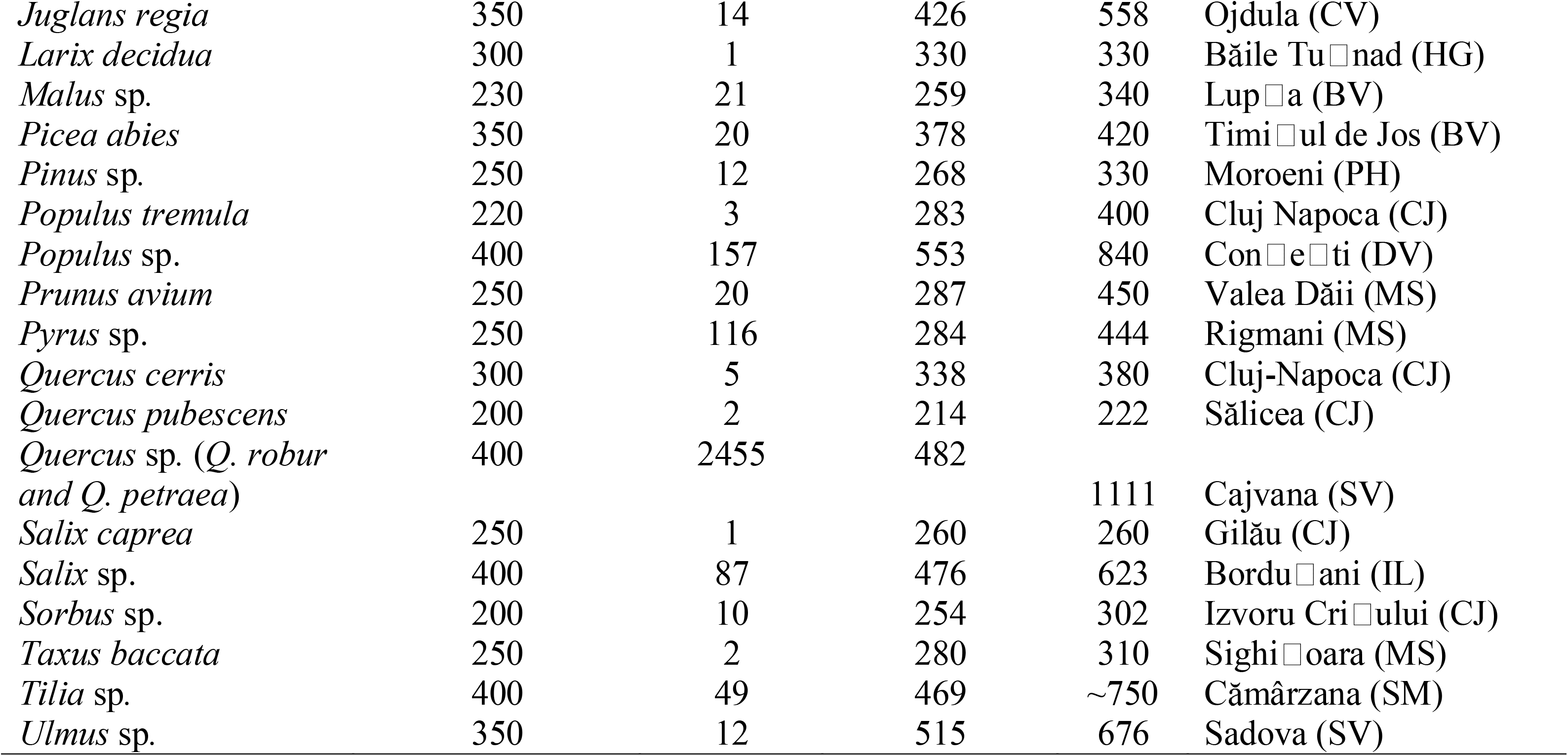
Descriptive statistics for the large old, single stemmed trees. The trunk circumference threshold represent the lowest size we considered for naming a tree ‘large old’

### Framework for understanding the distribution and conservation of large old trees in Romania

Figure 1 presents the framework we propose and employ to provide a comprehensive and structured understanding of the current knowledge and conservation of large old trees in Romania. Existing knowledge is reviewed on three main dimensions: large old trees, regulations relevant for large old trees and the ways how the large old trees are present in the local (regional) Romanian newspapers. Below we will present the ways we searched, analyzed and interpreted the informations on Romanian large old trees to achieve the objectives, with references to the framework presented in Figure 1.

**Figure 1.**
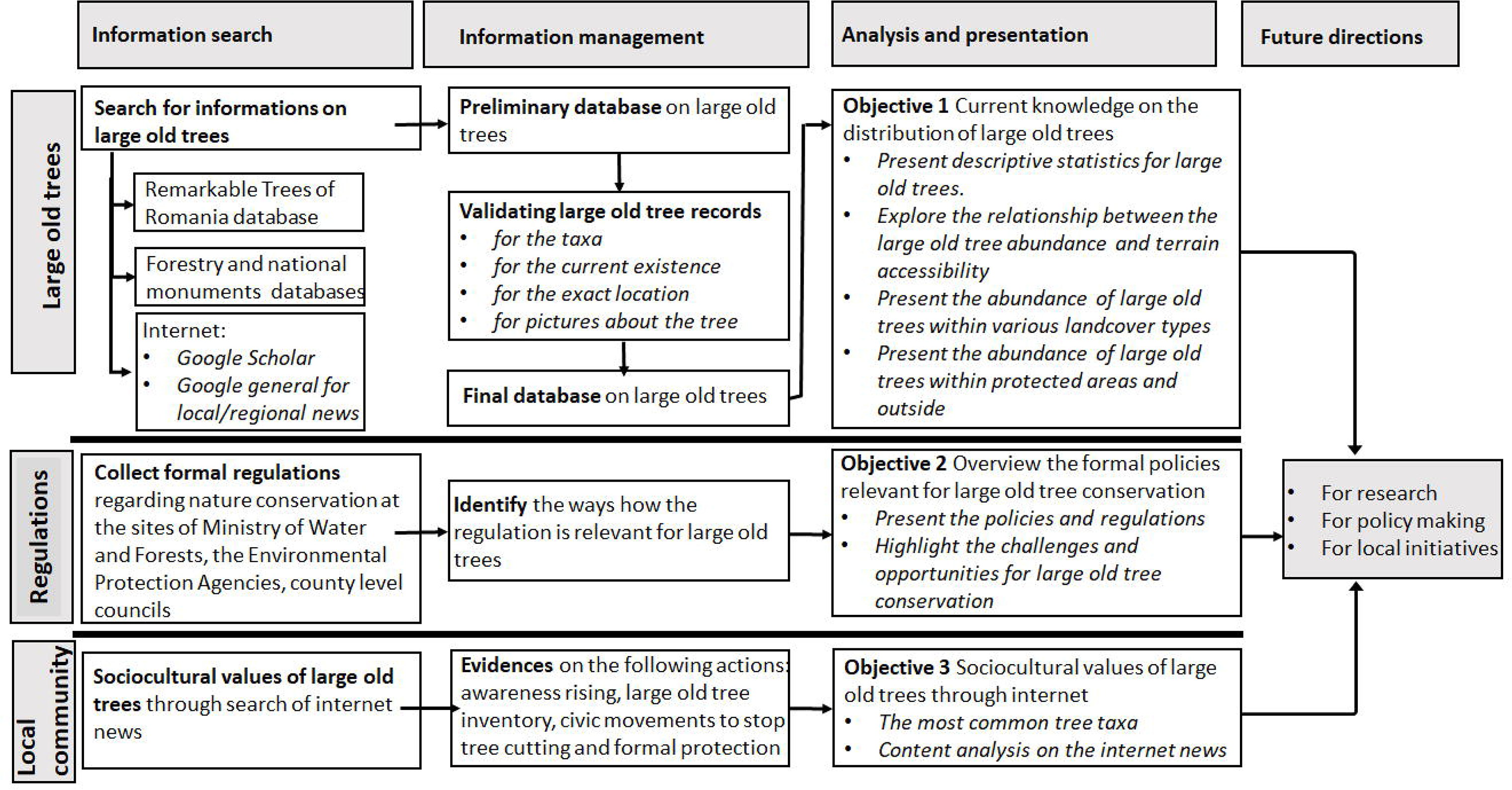
The framework used in this study.

### Information search and management

#### Large old trees

Currently the web platform Remarkable Trees of Romania (RToR) is the largest open access database for the Romanian large old trees (https://arboriremarcabili.ro/en/about-project), with over 4900 trees 4032 of them being native species. The platform was initiated in June 2015 in order to motivate large old tree inventory in the field, using as inspiration the well-established Ancient Tree Inventory of the Woodland Trust from UK. The majority of the large old trees (93%) on the RToR platform were uploaded by V. Arghius and the RToR team and were identified in the field. Second, we carefully searched articles from the journal of the Forestry Association ‘Progresul Silvic’ entitled ‘The Journal of Forestry and Hunting’ (‘Revista de Silvicultură□i Cinegetică), in order to identify other references to large old trees (Figure 1). Progresul Silvic often refers to ‘The National Register of Exceptional Trees in Romania’ when mentioning or presenting large old trees in Romania. Although the above register is not open for public, the large old trees presented in the Journal of Forestry and Hunting are referred as being present in the register (e.g., Radu and Coandă, 2006, Bolea, 2013a, Bolea, 2013b, Vasile et al., 2013, Lupu et al, 2015, Popa, 2016, Bolea, 2017). We also searched the formal List of Historical Monuments of Romania, available for every county at the webpage of the Romanian Ministry of Culture (2015, https://tinyurl.com/tuh65rs). In these lists every monument is exactly located (city, street number, name) and these informations together can be further used for internet search.

Third, we used specific keywords such as ‘monumental trees Romania’, ‘large old trees Romania’ to search in Google Scholar for academic publications on large old trees (Figure 1). Fourth, we searched on internet in Romanian and Hungarian languages (Hungarians being the largest ethnic minority in Romania) in order to identify informal publications (electronic news, informations available on YouTube channel) about the existence of large old trees (Figure 1). The keywords were ‘natural monuments’, ‘protected trees’, ‘old trees’, ‘large trees’, ‘secular trees’, ‘monumental trees’, ‘tree of the year’. These keywords were used alone as well as in combination with the county (regional administrative units) and with the most common and popular large old trees (oak, lime, elm, poplar, ash, beech, pear, fir). In the large majority of the cases, the different sources of informations (see above) validated each other (i.e. the same trees were present in multiple surces). Every large old tree identified in the internet search was ‘validated’ through checks for pictures, descriptions and locations with the use of satellite images available on the internet (e.g. Google Maps, Google Earth which sometimes allows even the determination of taxa via street view function, i.e. through tree leaves). For those trees which had exceptional sizes (DBH or CBH) we made in situ verification to confirm their existence, size and taxa. Every large old tree identified through internet search about which we had all the above mentioned informations was uploaded on the RToR site to provide the most updated online platform on large old trees from Romania. Only the native large old trees from this final database (at 15 January 2020) were analyzed for the current study (Figure 1).

#### Formal regulations relevant or potentially relevant for large old trees

Currently there are no formal regulations in Romania which explicitly targets the conservation of large old trees, regardless their position (i.e. protected areas or outside). Therefore we relied on the overview of the existing regulations targeting the nature conservation and environment which are also relevant for large old tree protection in Romania (Figure 1). This was done based on our direct experience as collaborators for developing management plans protected areas in Romania as well as through consulting other colleagues with relevant expertize on nature conservation legislation.

We also searched for regulations available at the webpages of the Ministry of Water and Forests as well as of the Environmental Protection Agencies (EPA). Furthermore we searched for formal decisions through which individual trees are protected at local (i.e. locality) and county levels. According to the Romanian formal procedures whenever natural elements are protected at local or county levels, the formal documents declaring their protection also enumerates the existing Laws and Ordinances based on which they are valid. After compiling the regulations relevant to large old tree conservation we analysed them in order to highlight how they are relevant to the conservation of large old trees and what further steps are necessary to further improve their relevance on this respect (Figure 1).

#### Large old trees and the local communities (electronic news search)

From an initial 550 internet links, containing electronic sources about the large old trees, we carefully removed the redundances (i.e. the same news about a specific tree is picked up by multiple platforms) and remained with 259 links for analysis.

Every electronic news was carefully read and grouped according the following 13 categories: *tree related* informations (if the news shared biological and geographical informations about the large old tree), *identity* (reference to the identity of the place and / or the local community), *history* (referring to historical informations about the place, for example unification of territories), *experience* (denoting personal and collective experiences related to large old trees), *revolt* (when local community members revolted against authorities which treatened the large old trees), *spiritual* (religious or mithological references), *gathering* (cultural gathering around the trees), *vandalized* (when the large old tree was injured by people), *tree of the year* (when the large old tree was nominated to national or international competition), *scared people* (when people from the local community were concerned about the fact that the tree may cause damages if it is broken), *protection* (when the article/news calls for the protection of large old trees), *arts* (the large old tree was subjected to artistic maniphestations) and *hazards* (the tree was damaged by severe whether conditions). We considered only the main types of values and connections presented in the title and the central message of the papers.

#### Data analysis and information presentation

The large old tree occurrence records compiled from various sources were used for statistical analyses (Figure 1). We created a map using 10×10 km grid cells where we show the distribution patterns of the validated native large old tree records in Romania. The map was generated using ArcGIS 10.3.1 software. The Romania teritory was divided into cells based on the official Romanian border in shapefile format (downloaded from the Romanian INSPIRE Geoportal http://geoportal.gov.ro/Geoportal_INIS/; coordinate system epsg:3844). In order to divide the teritory we used the Fishnet tool in the above mentioned software package. All resulting cells were assigned an unique numeric code. After this, we explored if the number of large old tree records within the 10×10 km cells is related to some broad accesibility features of the grid cell, such as: the depth of the fragmentation, the density of fragmentation, the average distance of the center of the cell from the nearest main road as well as the average elevation above sea level. These variables and the relationship between the large old trees and these variables are described in Annex 1.

We also present the number of large old tree records in the context of the current protected areas in Romania. For this we present informations on the total amount (km^2^) of land which is under any kind of nature protection regulation in Romania, the number of large old tree records which are within and outside protected areas and the total amount of protected areas (km^2^) from which no openly available large old tree records exist which could be validated (see above). We use *t-test* to compare the average CBH of the most common large old trees from our database (oak, beech) within the protected areas and outside.

In order to understand the patterns of co-variation within the data describing the large old trees in electronic newspapers we opted for Principal Component Analysis (PCA). In establishing the number of Principal Components, we visualized the correlation matrix between the variables with a network diagram (with *qqgraph*). This initial inspection highlighted that variables tend to form two clusters. We established the PC’s to 2 on the base of the eigenvalues (1 or more), the complexity values (ideally close to 1) and the uniqueness values of the variables on the PC. The PCA was run with the function *principal* (package *psych)* and specifying the *tetrachoric* (‘tet’) for the correlation type, our variables being binary. We removed variables which had very high proportion presence *(Tree,* 65%) as well as the very rare variables *(Scared, Arts<5%).* These analyses were implemented in R (version 3.6.0, 2019).

## Results

### The current knowledge on large old trees of Romania

There are 4032 validated native large old tree records in Romania (Table 1). From a total of 2544 10×10 km grid cells, large old trees were recorded in 311 (indicating 12% of the cells). The number of tree records per cell is positively skewed (*γ* = 7.14) and leptokurtic (*β* = 64.19), with few cells having high tree abundances of large old tree records, especially in the central and North-Western part of the country (Figure 2). The abundance of the large old tree records within the grid cells was significantly related to the depth and density of fragmentation and only marginally significantly with the average road distance, with significant quadratic effects for the first two variables (Table S1, Annex 1). Thus, the abundance of large old tree records increased with the increase of the fragmentation variables till a threshold was reached, after which (i.e. with the increase of terrain ruggedness and inaccessibility) the abundance of records sharply decreased within the grid cells with large old tree records (Figure S1, Annex 1). The average number of large old tree records peaked at 500 and 1200 m average altitudes, these being influenced by the abundance of the records of the two most common tree taxa (*Oak* and *Beech)* (Figure S2, Annex 1).

**Figure 2.**
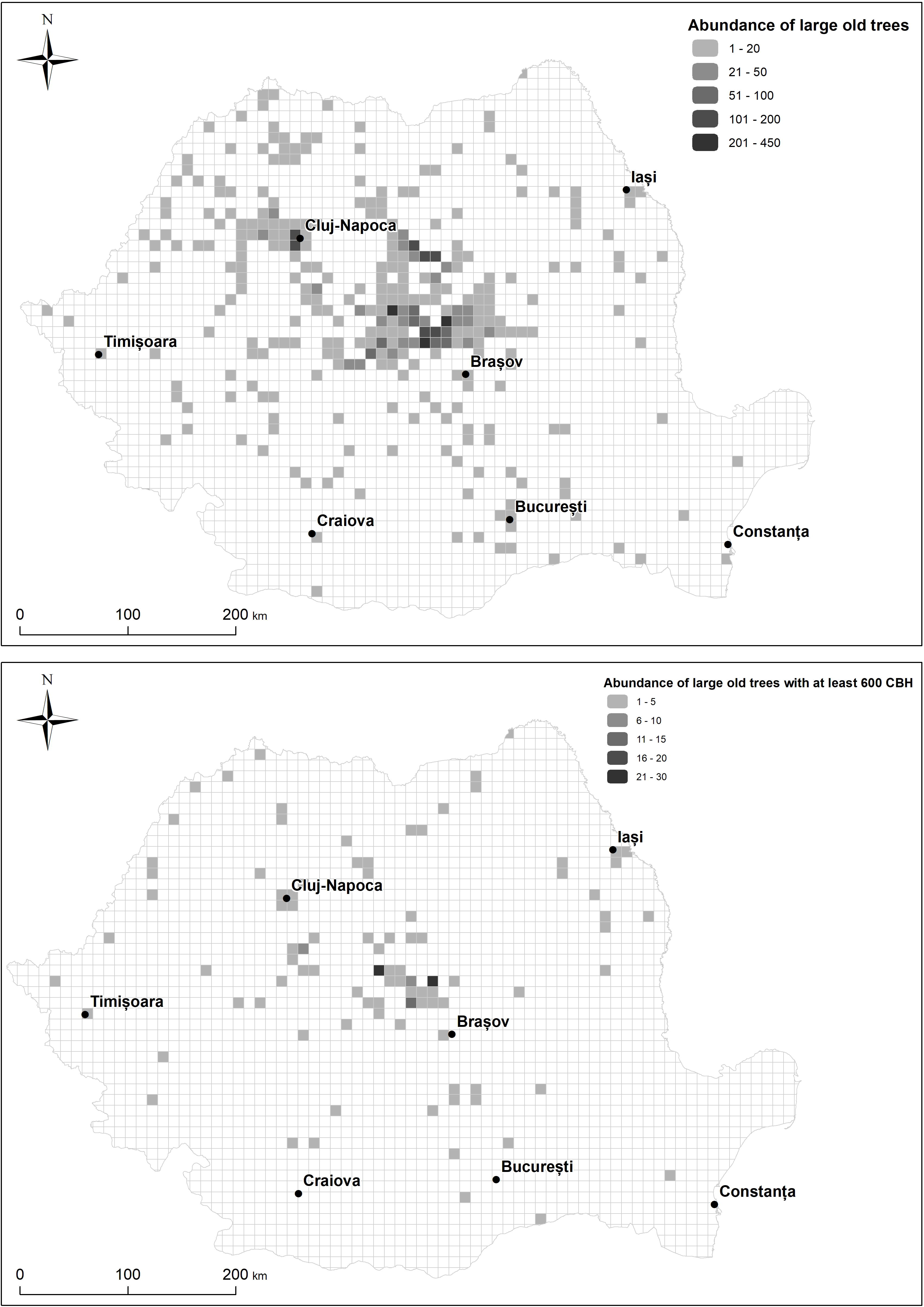
The abundance of large old tree records in Romania based on 10×10 grid cells.

The maximum tree sizes for the existing native large old tree taxa in Romania are presented in Table 1. The built areas, forests, meadows and pastures are the richest in tree taxa (Table 2). The *Oak, Poplar* and *Ash* are the most common trees in all landcovers excepting the urban parks where the *Ash* is dominating. The rarest trees are the *Chestnut, Service tree, Apple, Alder, Walnut, Elm, Birch* and *Fir* (Table 2).

**Table 2.** The number of large old trees, the richness of genera as well as the most common and rarest trees in eight landcover types

| Landcover | Number of taxa | Three most common genera (n) | Rarest genera (n) | Number of trees |
| --- | --- | --- | --- | --- |
| Arable fields | 11 | Poplar (26), Oak (24), Willow (13) | Chestnut (1), Hornbeam (1),<br>Service tree (1), Lime (2) | 83 |
| Built area | 20 | Oak (79), Poplar (34), Willow (29) | Apple (1), Chestnut (1), Pine (1),<br>Hornbeam (2), Yew (2) | 241 |
| Forest | 21 | Oak (306), Beech (144), Hornbeam<br>(32) | Ash (2), Elm (2), Service tree (2) | 611 |
| Mixed agricultural<br>lands | 14 | Oak (134), Poplar (14), Beech (16) | Ash (1), Service tree (1), Alder<br>(2) | 214 |
| Orchards and<br>vineyards | 5 | - | Walnut (3), Poplar (2), Willow<br>(2), Oak (1), Pear (1) | 9 |
| Parks (green<br>spaces in human<br>settlements) | 8 | Ash (18), Poplar (15) | Birch (1), Maple (1), Pine (1) | 46 |
| Pastures and<br>meadows | 19 | Oak (1906), Beech (686); Pear (98),<br>Hornbeam (40) | Elm (1), Fir (2), Walnut (2) | 2737 |
| Wetlands | 4 | Poplar (37) | Maple (1), Willow (3) | 47 |

There are 57471.01 km^2^ of protected areas in Romania. From a total of 4032 validated large old tree records 2296 (57%) were in protected areas and 1736 (43%) were out of protected areas (Figure 4). There are 37875.18 km^2^ (cca 66% of the total) protected areas in Romania from which we could not identify large old tree records. Referring to the most common large old trees, 61% of the oaks were in protected areas. The average CBH of the oaks within protected areas was 476.69 cm (SD=62.99) and outside the protected areas was 471.82 (SD=82.49), the difference being only marginally significant *(t-test,* P=0.10). 57% of the Beech trees were in protected areas. The average CBH of the Beech in protected areas was 410.01 cm (SD=53.12) and outside was 411.68 (55.90), the differences being not significant *(t-test,* P=0.66).

**Figure 3.**
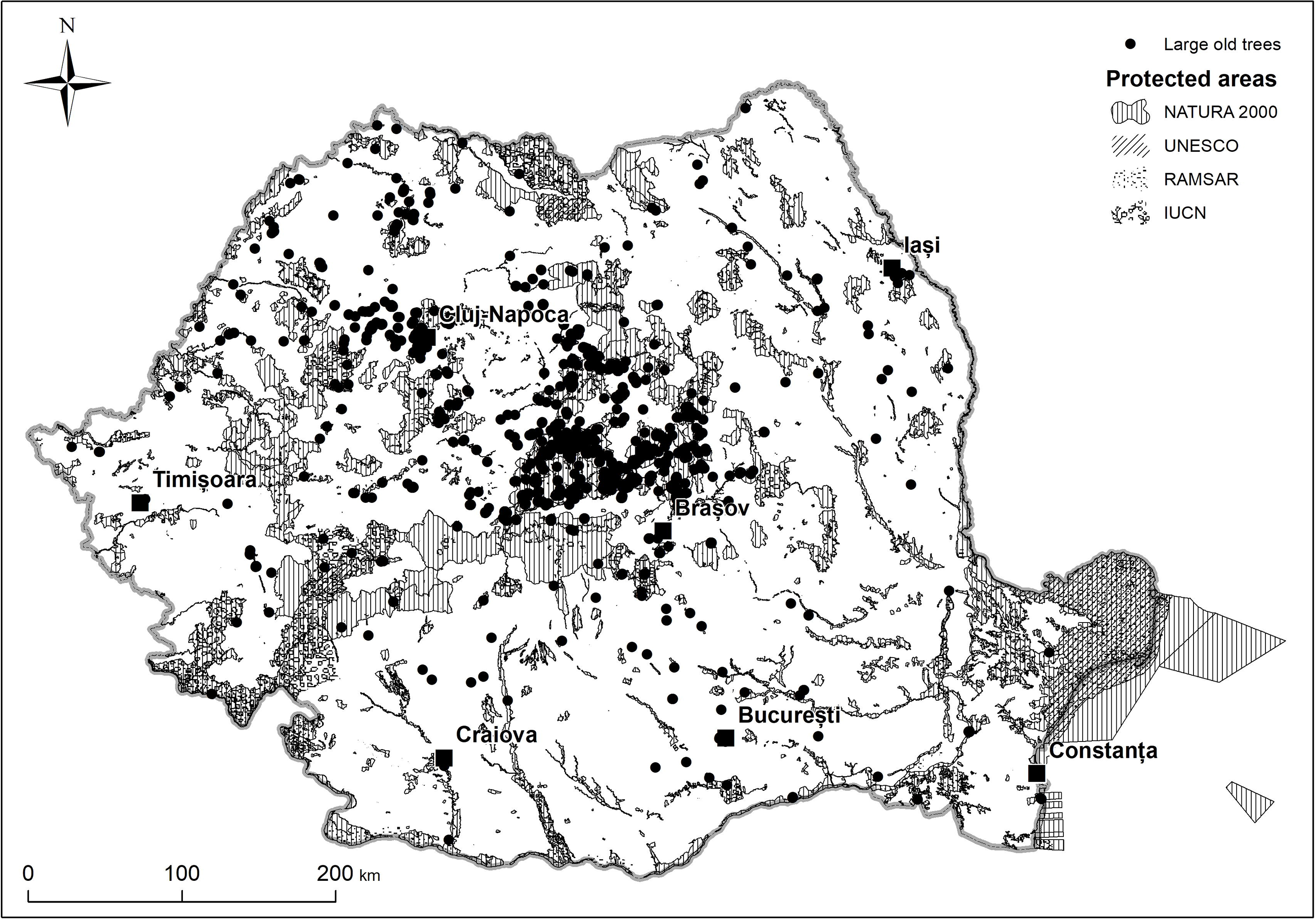
Map of the large old tree records and the protected areas in Romania.

**Figure 4.**
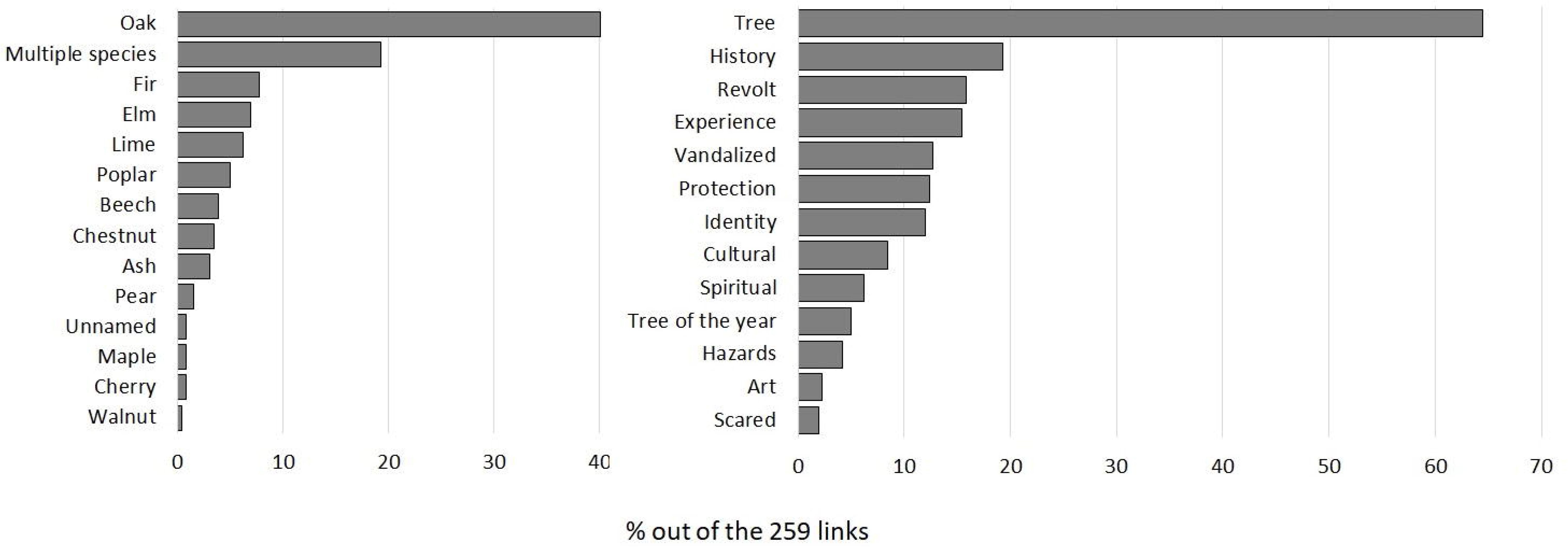
The proportion of internet news related to large old trees identified on electronic newspaper content analysis (taxa and types of informations).

### Formal regulations relevant to large old tree conservation

The Decree No. 237 from 1950 was the first – yet no longer valid – formal regulation which targeted natural monuments, including large old trees in Romania (Mohan and Ardelean, 1993b). Currently, there are two regulations which specifically target large old trees in Romania, while several other regulations can indirectly contribute to the conservation of large old trees in built areas as well as production landscapes if motivation and willingness exist at formal institutional levels to achieve their protection. Table 3 provides a brief overview of the current regulations existing in Romania regarding the protection of large old trees and it highlights the relevance, challenges and opportunities of those regulations for the protection of large old trees. The main challenges, regardless the regulation types, are related to the lack of robust expertize (arboriculture) and technology, the lack of knowledge and acknowledgement of the multiple sociocultural values of the large old trees at the level of the formal institutions. Furthermore, the regulations can often be misinterpreted and abused by the local authorities and landowners (Table 3).

**Table 3.**
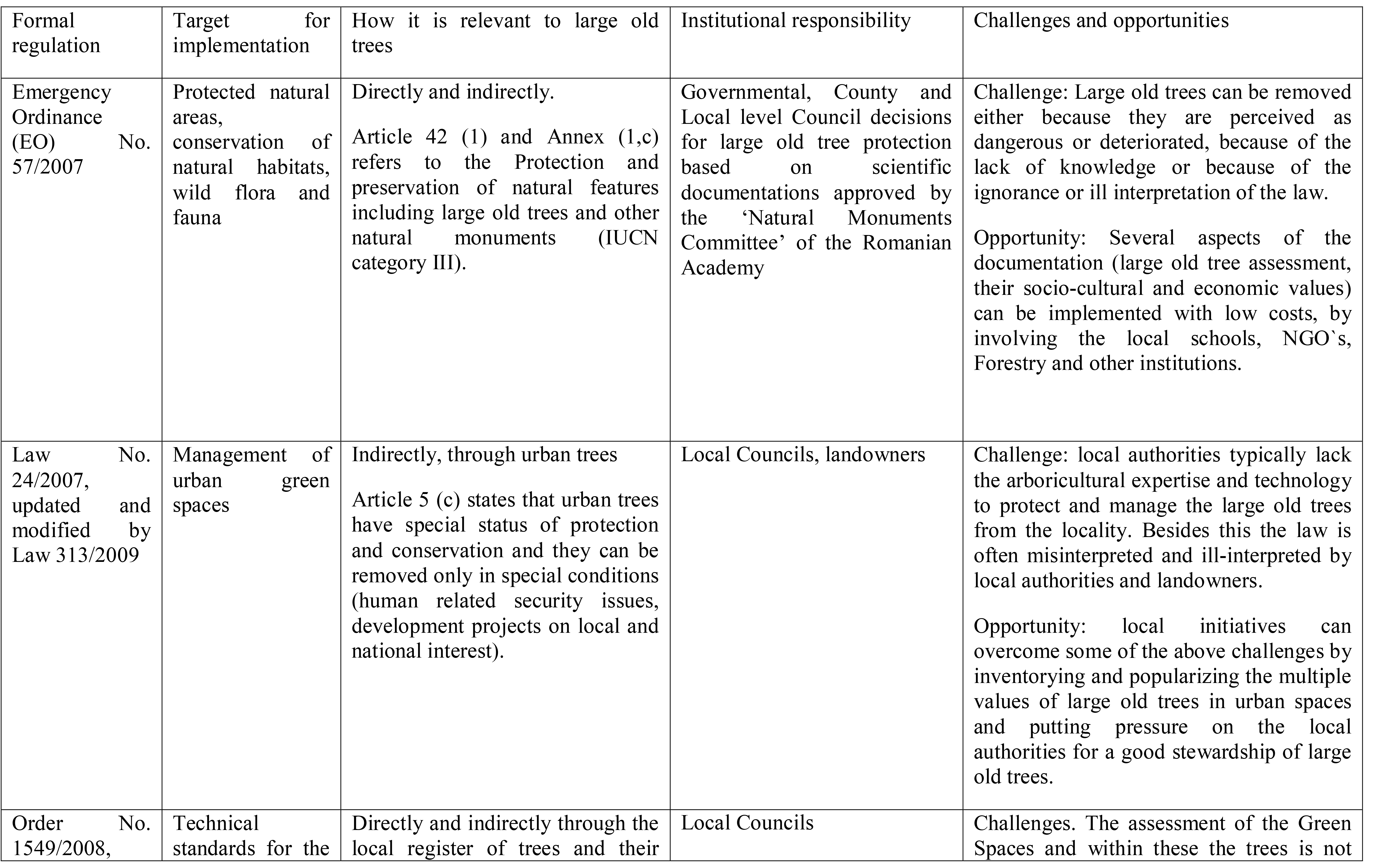
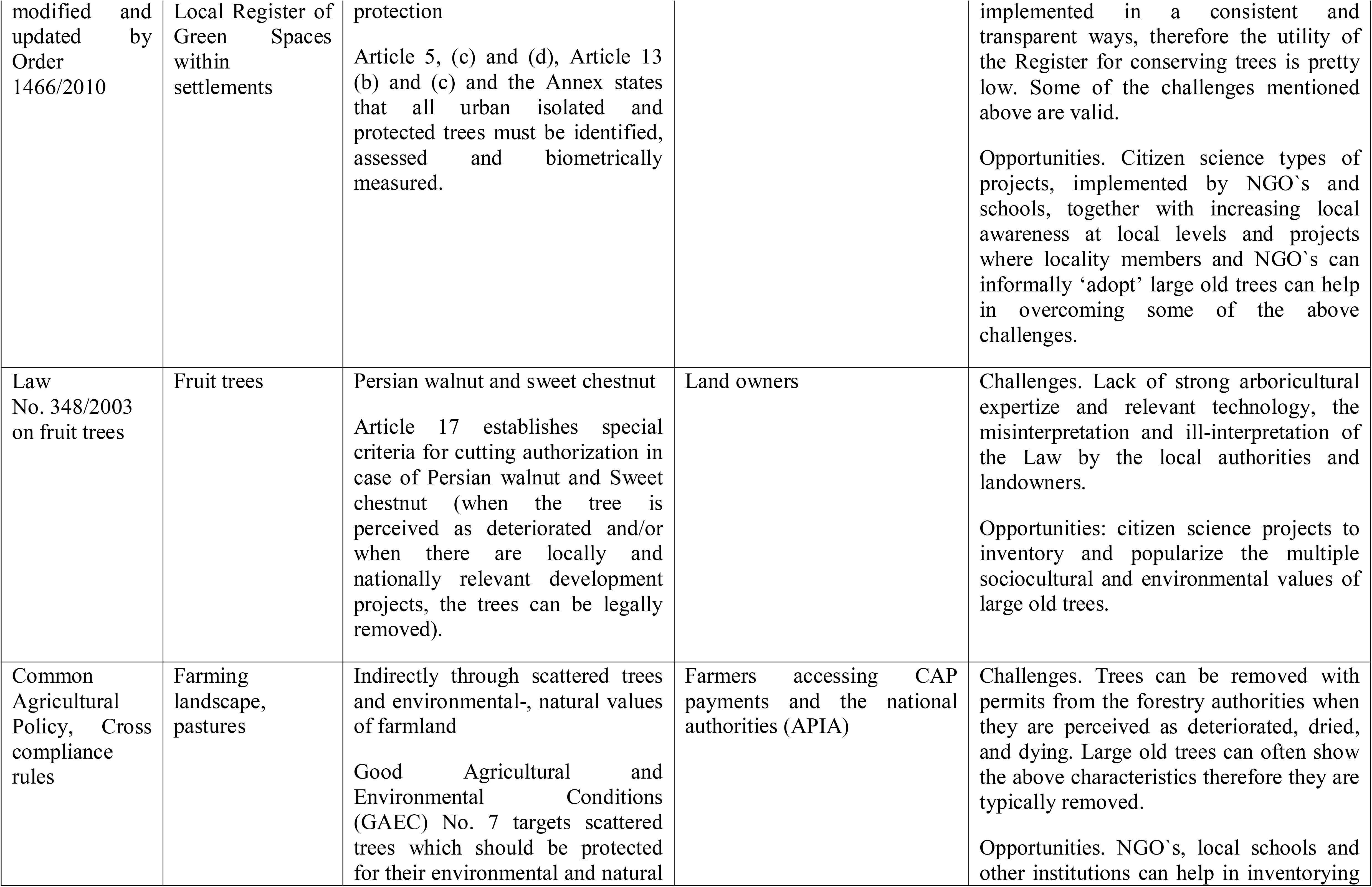
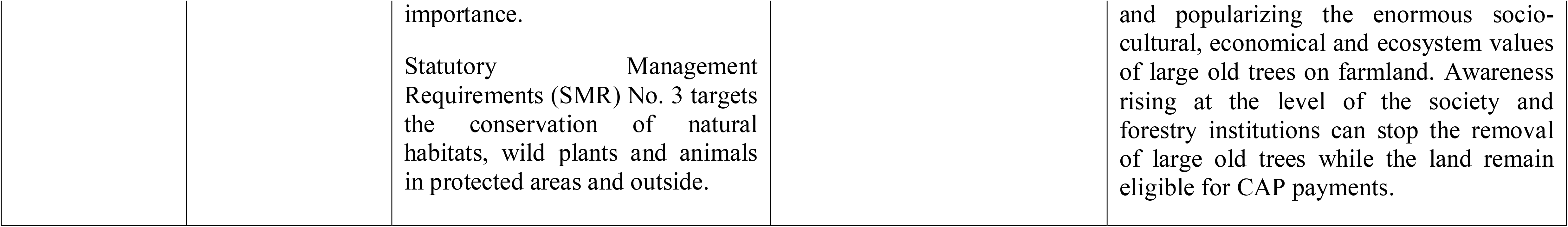
The formal regulations, their target and relevance for large old trees in Romania

There are several opportunities for recognizing and protecting large old trees at the levels of the local communities, including citizen science types of projects for inventorying and documenting the socio-cultural values of large old trees at local levels, awareness rising activities and assisting local authorities to recognize and protect large old trees (Table 3). Several of these activities are low-cost and can be implemented within educational collaborative projects between local schools and local NGO’s. For example in the town of Sighisoara, the collaboration between the local schools and the Association Eco-Breite resulted in the inventory of over 400 large old oaks in the Breite ancient wood-pasture nature reserve and these informations were used for the formal extension of the protected area to capture large old trees which were not protected by nature conservation legislation (T. Hartel was part of the team as main organizer of the school activities). The Association Kogayon organized in 2014-2016 a citizen based inventory project of large old trees entitled ‘The Grandparents of the Forest’ in the National Park Buila-Vânturari □ a and its surroundings. There were over 250 large old trees inventoried and documented in order to develop conservation strategy for these trees (source of informations: Facebook page of Kogayon Association: https://www.facebook.com/search/top/?q=bunicii#20padurii%epa=SEARCH_BOX). Large old trees from the county of Ia□i were inventoried and formally protected as an initiative of the Environmental Protection Agency and the County Level Council (http://www.icc.ro/sites/default/files/files/online/noutati/Arbori.pdf).

### Large old trees in the Romanian newspapers

Following the electronical news searches about the large old trees of Romania references for 12 tree taxa have been identified. In some situations the news were related to multiple or unidentified species. The *Oak* was the most popular taxa, being present in 40% of the sources, the multiple species large old tree category was the second most frequent (cca 20%) while the *Maple, Cherry* and *Walnut* were the rarest (<1%) (Figure 4). Regarding the information topic of the newspapers, cca 65% of these targeted the description of the trees (*trees*) (65%) and the rarest were about the *tree of the year* competition, natural *hazards* affecting the trees, *arts* and *scared people* (Figure 5). We also highlight the relatively high proportion of the information regarding *revolt* (see above, 16%, Figure 4) and the need to *protect* these trees (15%, Figure 4).

The PCA analysis of the newspaper information content on large old trees shows that the PC1 was characterized by high loadings for the variables *identity, history, gatherings* (positive), *revolt* and *vandalized* (negative) (Table 4). Based on these variables, the identity of the PC1 reflects a deep social concern towards the fate of the large old trees in their community and the appreciation of the large old trees for their socio-cultural values. PC2 was characterized by high loadings of the variables *spiritual* and *hazards* (both positive) (Table 4). This suggests that these reports also highlights the negative impact of the unusual weather conditions on the large old trees.

**Table 4.** The results of PCA on the key topics present in the popular news on large old trees, identified on the internet.

| Variable | PC1 | PC2 | Uniqueness | Complexity |
| --- | --- | --- | --- | --- |
| Identity | **0.75** | -0.21 | 0.40 | 1.2 |
| History | **0.78** | -0.05 | 0.39 | 1.0 |
| Experiential | 0.40 | -0.38 | 0.69 | 2.0 |
| Revolt | **-0.85** | -0.28 | 0.19 | 1.2 |
| Spiritual | -0.13 | **0.83** | 0.08 | 1.7 |
| Gatherings (cultural) | **0.62** | -0.25 | 0.55 | 1.3 |
| Vandalized | **-0.89** | -0.29 | 0.13 | 1.2 |
| Tree of the year | 0.10 | 0.32 | 0.89 | 1.2 |
| Scare people | 0.04 | 0.36 | 0.87 | 1.0 |
| Protection | 0.11 | -0.02 | 0.99 | 1.1 |
| Hazards | -0.18 | **0.78** | 0.36 | 1.1 |
| SS loadings<br>(eigenvalue) | 3.31 | 1.84 |  |  |
| Cumulative variance | 0.30 | 0.47 |  |  |

The mean item complexity was 1.2 suggesting that the variables tend to be well represented in one PC (R-bloggers, 2018, https://tinyurl.com/vxhalvg, Table 4). The uniqueness representing the proportion of variance which is unique for a specific variable also highlight that the above highlighted variables have high relevance for the given PC’s (R-bloggers, 2018, https://tinyurl.com/vxhalvg, Table 4).

## Discussions

To the best of our knowledge, this is the first national holistic assessment of large old trees. Our framework simultaneously considers the informations related to the large old trees (taxa, sizes, current knowledge on their distribution), it provides a critical assessment of the existing regulations with opportunities for protecting large old trees at the level of localities and counties, and a comprehensive understanding of the information types currently existing in the electronic news in the addressed country (Romania). Our research is timely because the great majority of the countries in Europe and worldwide still don’t reported the situation of their large old trees in academic journals and our framework can be used as a guide for a holistic assessment and reporting of these trees. Our framework can be easily implemented because it largely relies on data compilations and internet search. Furthermore, the holistic perspective can contribute with important case studies to the emerging new concepts for sustainability and biodiversity conservation research, including the ‘leverage points’ (Abson et al., 2016) and ‘seeds of a Good Anthropocene’ (Bennett et al., 2015).

### Knowledge and distribution of large old trees

Our results suggests that there in are large areas in Romania with no or very few large old tree data available; notably continuous 10*10 km cells with large old tree records exist only in the Central and the North-Western parts of Romania (Figure 2). Furthermore we found a quadratic relationship between the terrain accessibility variables and the abundance of the large old tree records within the cells (Annex 1, Figure S1), indicating that increasing inaccessibility the abundance of large old trees in the cells is decreasing. This shows that within the cells where large old trees are recorded, the search tends to be opportunistic, i.e. the easily accessible parts of the landscape are more likely to be surveyed. Furthermore, the research projects targeting ancient wood-pastures also targeted elevations where the broadleaf trees dominated (Oak-Beech) and not the mountain areas (Figure S1, Annex 1 this manuscript Hartel et al., 2018). Despite the lack of an even coverage of large old tree records in Romania, biologically and ecologically reliable informations can be extracted from the current data. For example the most common large old trees in the arable fields are the Poplar, Oak and Willow (Table 2). Poplars and willows are known for their relatively high soil humidity reqirements (Lindroth et al., 1994; Petzold et al., 2011), several of these trees are distributed along the past flooded areas of the rivers (which are now used as arable fields). Nevertleless old willows in arable lands are typically pollarded, suggesting that these trees had a key role in the local communities (by providing woody material). The large old oak trees in the arable fields may be also legacies of the past landuses, specifically the grasslands which were used as pastures or hay meadows and were converted into more intense arable lands (Plieninger et al., 2015). Forests are dominated by the oak, beech and hornbeam, which can be expected considering the dominant tree species in this land cover (Doni□ă et al., 1992). Several large old oak trees in the hornbeam dominated forests of Central Romania are the legacies of the past wood-pastures which were reforested in the past decades after the collapse of the communism. In the Natura 2000 site Sighisoara-Tarnava Mare for example there are over 20,000 hectares of closed forest which formally are still pastures. In these forests the abundance of open grown large old oak trees is high (Hartel et al. 2018). Large old oak trees of which shape suggests that the tree was open grown (i.e. large, rounded crown) in forests can help in a better understanding the history of that particular forest (Kirby, 2014).

We found that near half of the existing and validated large old tree records in Romania are from areas which have no formal conservation protected status. Furthermore, over half of the protected terrestrial area surfaces of Romania have no openly available large old tree records and based on the two most common taxa, protected areas does not protect larger trees than the unprotected areas. An accumulating number of evidences in Europe shows that large old trees have exceptional ecological values, serving as habitats for a large number of protected species in Europe in open pastures, parklands, forests, orchards and urbanized areas (see e.g. Horak, 2014; Horák and Rébl, 2012; Koch et al., 2012; Lachat et al., 2012; Miklín et al., 2017; Müller et al., 2014; Sebek et al., 2013). As the protected areas in Romania were classically underfunded and suffers from lack of reliable informations regarding the biodiversity based on which management actions can be implemented (Ioja et al., 2010) citizen science types of activities could help in inventorying large old trees (Loos et al., 2015). We mentioned two citizen based activities to inventory and protect large old trees in protected areas of Romania (see Results) and expressions to protect these trees are also present in the Romanian internet news (see below). We do not exclude that large old tree inventory and protection initiatives in Romanian protected areas may be more common than we show based on our validated large old tree data, these activities being not available to the broad public cannot represent actionable knowledge based on which similar initiatives or the conservation of the inventoried trees can be implemented.

### Formal regulations as challenges and opportunities for large old tree conservation

In this overview we also present the current context of formal legislation in Romania which is, or may be relevant for the protection of large old trees. Within the current legislation of Romania we identified five regulations which are relevant to large old tree conservation and from these, only two are directly relevant to large old trees: one targeting the protected areas and one targeting the green spaces within settlements (Table 3). Besides the lack of mentioning explicitly the large old trees, the challenges related to the implementation of the regulations in the benefit of large old trees are related to the lack of knowledge (intent), the lack of expertize, institutional capacity, vested interests (corruption) and inconsistencies within the regulations. Several of these aspects are also key challenges for implementing existing regulations regarding nature conservation (Hossu et al., 2017; Ioja et al., 2010), agri-encironmental payments (Mikulcak et al., 2013), rural- (Mikulcak et al., 2015), and urban development (Grădinaru et al., 2020). Nevertheless, even with the above challenges, the current legislation does allow the inventory and protection of the large old trees if this is locally desired. We highlighted as example the county of Iasi, where for the first time in Romania, every large old tree is formally protected (see Results), although the raw data (e.g. CBH) are not publicly available about these trees. Local initiatives can also save large old trees. An eloquent example in this respect comes from the commune of Saschiz, where a research on large old trees on wood-pastures highlighted several local stories about a specific large old tree, known especially by the elderly people (Hartel, 2008). These stories were collected and shared on social media platforms (facebook, T. Hartel), and served as motivation for local initiatives to identify the tree and include it into the touristic destinations of the commune (Hartel, 2008). Furthermore, Moga et al. (2016) presents the case of the second largest oak from Romania, which was discovered by an educational activity and then formally protected, being now part of the cultural identity of the commune of Mercheasa (Box 1 in Moga et al., 2016).

### Large old trees in the Romanian news

We collected and analysed the available informations on large old trees of Romania from the internet. The non-academic types of informations available on the internet such as those published on the local newspapers and Youtube movies highlights the ways how the local communities and/or their leaders potentially perceove the presence of large old trees in their surrondings and the condition of these trees. We showed that there are a large number of tree taxa present in the electronic news, the oak being the most common. Furthermore, we show a wide range of values and concerns associated to large old trees at the level of the local communities. Informations about the large old tree (taxa and/or size and/or origins and/or age) were the most common, as expected. Furthermore, several values which are reflecting the importance of large old trees for the local community in a relational sense (sensu Chan et al., 2016) were expressed, including the historical-, identity-, cultural and spiritual values of the large old trees as well as the concerns related to their human induced deterioration and revolts at the level of the local community against formal decisions to cut the trees. Furthermore a genuine expression exist to protect these trees. In combination with the above presented legislative context, we see that desire exist at the level of the local community to value, protect and formally include the large old trees in the historical and monumental identity of the loality, however, we also perceive a huge gap between the institutional effectiveness to build on these intentions and values. The outstanding sociocultural values of the large old trees were shown in recent overviews (Blicharska and Mikusiński, 2014; Hartel, 2018) and suggests that exploring the values existing at the leven of the local communities can guide conservation actions for large old trees.

## Conclusions: implications for science, policy and local initiatives

With this analysis we aim to provide a holistic to understand and address the conservation challenges of large old trees, by simultaneously considering every available information on the distribution of these trees, presenting the formal regulations directly or indirectly relevant to large old trees and by providing a comprehensive analysis of the current knowledge on large old trees as present in the internet news. Holistic approaches such as that proposed by us can guide the conceptual framing of research related to large old trees: being old and often having large size, these trees have several social and ecological values, which can be simultaneously addressed using integrative conceptual frameworks. Recently proposed sustainability concepts to which maturation and contextualization such assessments could contribute are ‘leverage points’ (Abson et al., 2016; Fischer and Riechers, 2019) and the ‘seeds of a good Anthropocene’ (Bennett et al., 2015). Both concepts considers the importance of human agency in shaping the present and future of the various components of nature or the environment and the potential of social transformations to protect the natural environment. Conceptual development can help the development of better policies to protect large old trees. By a detailed overview of the Romanian legislation we found that large old trees are only scarcely addressed and formal instituions are generally too weak to effectively protect these trees. However, we also found that the society is genuinely valuing large old trees and this can be a huge opportunity for the decision makers to protect these trees. We provided recent examples for local and county level activities which resulted in the protection of large old trees highlighting the power of intent in achieving large old tree conservation. Local initiatives can contribute to the inventory, popularization and protection of the large old trees. While the civil societies are actively engaged in large old tree inventory and popularization in Western Europe, we only found few initiatives where local NGO’s mobilized local communities for large old tree protection (see e.g. Box 1 in Moga et al., 2016, Hartel, 2018, this paper). Our internet news research suggests that people may be open for such activities and they are concerned about the fate of their large old trees. Finally, we encourage every country to create open access web platforms and upload there the large old trees. Such platforms with open access tree informations help further and more complex social-ecological analyses related to the distribution and status of these trees.

## Acknowledgements

We are very grateful for Sir Charles Burrell for supporting the creation and maintenance of the web platform Remarkable Trees of Romania. We express our honest appreciation to Jill Butler, Isabella Tree and Ted Green for their encouragement and efforts to understand and protect Romanian large old trees and ancient wood-pastures.

